# STING deficiency-associated aberrant CXCL10 expression contributes to pathogenesis of arthritogenic alphaviruses

**DOI:** 10.1101/2020.05.13.095083

**Authors:** Tao Lin, Tingting Geng, Andrew Harrison, Duomeng Yang, Anthony T. Vella, Erol Fikrig, Penghua Wang

**Affiliations:** Department of Immunology, School of Medicine, University of Connecticut Health Center, Farmington, CT 06030, USA; Section of Infectious Diseases, School of Medicine, Yale University, New Haven, CT06520, USA

**Keywords:** STING, CXCL10, alphavirus, Chikungunya, viral arthritis

## Abstract

Arthritogenic alphaviruses such as Chikungunya virus and O’nyong nyong virus cause acute and chronic crippling arthralgia associated with inflammatory immune responses. However, the physiological functions of individual immune signaling pathways in the pathogenesis of alphaviral arthritis remain poorly understood. Here we report that a deficiency in the stimulator-of-interferon-genes (STING) led to enhanced viral loads, exacerbated inflammation and selectively elevated expression of CXCL10, a chemoattractant for monocytes/macrophages/T cells, in mouse feet. *Cxcl10*^*-/-*^ mice had the same viremia as wild-type animals, but fewer immune infiltrates and lower viral loads in footpads at the peak of arthritic disease (6-8 days post infection). Macrophages constituted the largest immune cell population in footpads following infection, which were significantly reduced in *Cxcl10*^*-/-*^ mice. The viral RNA loads in neutrophils and macrophages were reduced in *Cxcl10*^*-/-*^ compared to wild-type mice. In summary, our results demonstrate that STING signaling represses, while CXCL10 signaling promotes, pathogenesis of alphaviral disease.

## Introduction

Alphaviruses are a genus of single-stranded positive sense RNA viruses within the *Togaviridae* family. These viruses are mainly transmitted by mosquitoes and pose a public health threat worldwide, particularly in tropical/subtropical regions. Many alphaviruses are arthritogenic, including Chikungunya (CHIKV), O’nyong-nyong (ONNV) and Ross River viruses (RRV) *etc*. CHIKV is the causative agent of acute and chronic crippling arthralgia that was initially identified in Tanzania in 1952 ^1^. Since then, several major epidemics have been recorded on the Indian Ocean islands, India, Southeast Asia, which resulted in over 6 million cases ^2^. In late 2013, CHIKV emerged on the Caribbean islands, and has now spread to more than 50 countries across the Central and South America, including autochthonous infections in the United States and caused over 2.5 million infection cases (Sources: Pan America Health Organization). Approximately 50% of CHIKV-infected patients suffer from rheumatic manifestations that last 6 months to years, with ∼5% of the victims having rheumatoid arthritis-like illnesses ^3,4^.

During the acute phase of infection in humans (∼ two weeks), CHIKV infects many organs and cell types ^2^, induces apoptosis and direct tissue damage ^5-7^. The acute phase is also characteristic of robust innate immune responses, including high levels of type I IFNs, proinflammatory cytokines/chemokines and growth factors ^5,8-12^. Immune cell infiltration is a hallmark of acute CHIKV infection, including primarily macrophages, monocytes, but also neutrophils, dendritic cells, NK cells and lymphocytes ^2^. In the chronic phase, CHIKV arthritis may progress without active viral replication, typified by elevated expression of cytokines and immune cell infiltration ^2,13^. In particular, human arthritic disease severity is associated with a high level of serum chemoattractants for monocytes/macrophages/T cells, CXCL10 and CXCL9 ^14^. In mice, CHIKV infection leads to a low viremia lasting usually 5-7 days, which is limited by type I IFNs ^11,15-17^ and is subsequently cleared by virus-specific antibody responses ^18-25^. When inoculated directly into a mouse foot pad, CHIKV elicits overt arthritic symptoms including the first peak of foot swelling characteristic of edema occurring 2-3 days post infection and a second peak at 6-8 days post infection ^26^ with massive infiltration of immune cells into infected feet ^16,27-32^.

The stimulator of interferon genes (STING) participates in innate immunity to both DNA and RNA viruses. During DNA virus infections, STING signaling induces type I interferons (IFNs) after being engaged by a second messenger cyclic GMP-AMP (cGAMP), which is synthesized by a viral DNA receptor, cGAMP synthase (cGAS) ^33-38 39 40^. Our study with West Nile virus (WNV) and other recent studies conclusively demonstrate that STING is also important for the control of RNA virus infection in mouse models ^33,41-43^; likely through upregulation of type I IFNs ^33 44 45^, induction of a specific set of chemokines via Signal Transducer and Activator of Transcription 6 (STAT6) ^43^, translation inhibition of viral gene expression ^46^ and/or other unknown mechanisms. Intriguingly, STING signaling was recently shown to induce expression of negative regulators of innate immune signaling, such as suppressor of cytokine signaling (SOCS), to control Toll-like receptor (TLR)-mediated hyper inflammatory immune responses in a lupus mouse model ^47^. We herein report that a deficiency in the stimulator-of-interferon-genes (STING) led to enhanced viral loads, exacerbated inflammation and selectively elevated expression of CXCL10 in mouse feet. *Cxcl10*^*-/-*^ mice had the same viremia as wild-type animals, but fewer immune infiltrates and lower viral loads in footpads at the peak of arthritic disease (days 6-8 post infection). Macrophages constituted the largest immune cell population in footpads following infection, which were significantly reduced in *Cxcl10*^*-/-*^ mice. The viral RNA loads in neutrophils and macrophages were also reduced in *Cxcl10*^*-/-*^ compared to wild-type mice.

## Results

### A STING deficiency leads to enhanced viral loads, exacerbated inflammation and selectively elevated expression of CXCL10

STING signaling is essential for induction of immune responses to DNA viruses. Our studies with West Nile virus (WNV) and other recent studies conclusively demonstrated that STING is also important for the control of RNA virus infection in mouse models ^33,41-43^. This is also observed with CHIKV infection (**Fig. 1**). When inoculated directly into a mouse footpad, CHIKV elicits a brief viremia that lasts ∼ 5 days and overt arthritic symptoms including the first peak of foot swelling characteristic of edema occurring 2-3 days post infection and a second peak at 6-8 days post infection ^26^ with massive infiltration of immune cells into infected feet ^16,27-32^. Sting-deficient mice (*Sting*^gt/gt^) ^48^ presented a significant increase in viremia from 12hrs through 96hrs post infection (p.i.) compared to wild-type (WT mice) (**Fig.1 A, B**). Furthermore, the viral loads in the spleen were much higher in *Sting*^gt/gt^ than WT mice too (**Fig.1 C**). The viral loads in the feet were elevated at day 4 p.i. in *Sting*^gt/gt^ compared to WT mice; however, this difference disappeared by day 6 and 8 p.i. (**Fig.1 D, E**). *Sting*^gt/gt^ mice showed much greater footpad swelling from days 4 through 8 p.i., than WT mice (**Fig.1 F**). Histopathology analysis by Haemotoxylin and Eosin (H&E) staining demonstrated a significant increase of immune cells in muscle/synovial cavity/tendon in *Sting*^gt/gt^ compared to WT mice (**Fig.1 G, H**). Interestingly, although the viral loads in *Sting*^gt/gt^ joints were the same as WT at days 6 and 8 after infection (**Fig.1 D**), the joint disease manifestations were much more severe in *Sting*^gt/gt^ (**Fig.1 F-H**).

**Fig. 1.**
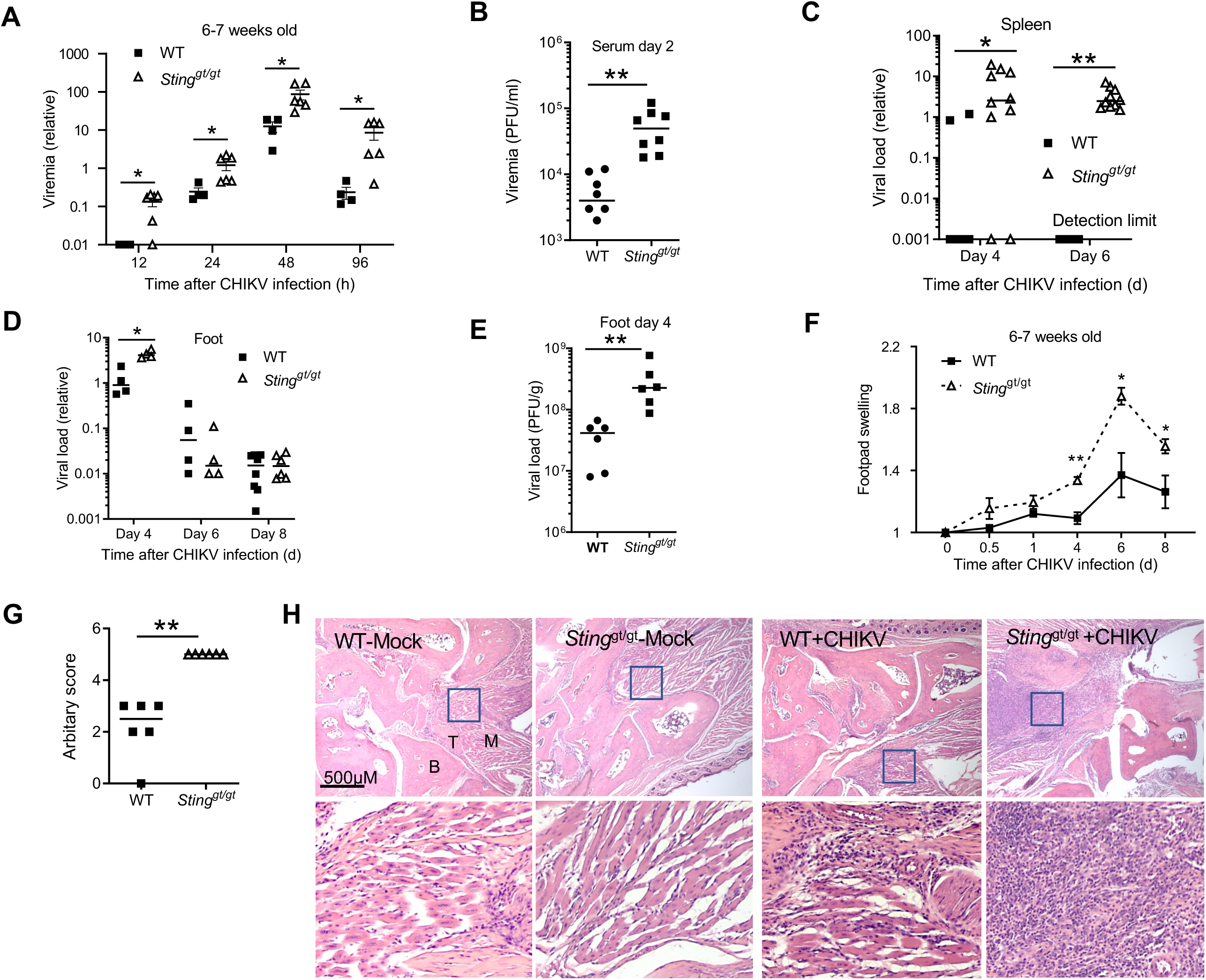
CHIKV pathogenesis is exacerbated in *Sting*^gt/gt^ mice. Age- and sex-matched, C57BL/6 (WT) and STING deficient (*Sting*^gt/gt^) mice were infected with CHIKV. **A**) quantitative RT-PCR (qPCR) quantification of CHIKV loads in whole blood cells. **B**) Viremia of day 2 sera quantified by plaque forming assay (PFU/ml serum). qPCR analysis of CHIKV loads in **C**) the spleen and **D**) ankle joints. **E**) Viral titers in footpads on day 4 pi (PFU/gram tissue). **F**) Fold changes in the footpad dimensions of infected (days 0.5, 1, 4, 6 and 8) over the uninfected (day 0) (N=4-6 per experimental group). **G**) Arbitrary scores of day 8 ankle joint inflammation and damage using a scale 1 to 5, with 5 representing the worst disease presentation. Each dot represents one mouse; the small horizontal line indicates the median of the result. N=6/genotype. **H**) Representative H&E micrographs of footpad inflammation 8 days after infection. N=6/genotype. Magnifications 40x. B: bone, T: tendon, M: muscle. Boxed areas indicate the regions with severe infiltration and tissue damage. Magnification: 200x. The data represent two independent experiments. *, P<0.05; **, p<0.01; ***, P<0.001 [**A-E**) non-parametric Mann-Whitney U test, **F** and **G**)] two-tailed Student’s t-test].

Since immune cell infiltration into joints and muscles is a hallmark of CHIKV arthritis, we examined chemoattractant expression in mouse ankle joints. A chemokine PCR array was performed with joint samples at the peak of disease (day 8 p.i). Two genes *Ifng* and *Cxcl10* (also known as IFN-γ inducible gene-IP10) were upregulated by 4-5 fold and three genes (*Cxcl5, Cxcr2, Ppbp*) were down-regulated by over 3 fold in *Sting*^gt/gt^ compared to WT mice (**Fig.2 A**). Downregulation of *Cxcl5* is consistent with upregulation of IFN-γ, as Cxcl5 can be inhibited by IFN-γ ^49^. Cxcl10 is a chemoattractant for T cells, monocytes/macrophages, natural killer cells (NKs), and dendritic cells (DCs); Cxcl5, Cxcr2, and Ppbp (also known as Cxcl7) participate in neutrophil recruitment. We next validated the PCR array results and investigated the kinetics of chemokine expression during the course of CHIKV infection. In WT mouse joints, *Ifng* and *Cxcl10* mRNA expression was continuously upregulated by ∼7-10-fold, and further elevated in *Sting*^gt/gt^ mice at days 4 through 8 p.i. (**Fig.2 B**), coincidently with arthritis progression. *Ppbp* expression was equally upregulated in both WT and *Sting*^gt/gt^ mice at day 4 p.i., but rapidly decreased at day 8 p.i. in *Sting*^gt/gt^ mice (**Fig.2 B**). Ccl5 (a chemoattractant for T cells, eosinophils and basophils) and Ccl1 (chemoattractant for monocytes), Ccl20 (strongly chemotactic for lymphocytes), Ccl26 (chemotactic for eosinophils and basophils) were transiently up-regulated at day 4 p.i. and down-regulated at day 8 p.i. similarly in both WT and *Sting*^gt/gt^ mice (**Fig.2 B**). These data demonstrate that IFN-γ and CXCL10 expression kinetics are in line with arthritis progression and suggest a role for IFN-γ and CXCL10 in disease pathogenesis. However, IFN-γ has been known to play an anti-CHIKV role ^50^ and thus is excluded for further investigation.

**Fig. 2.**
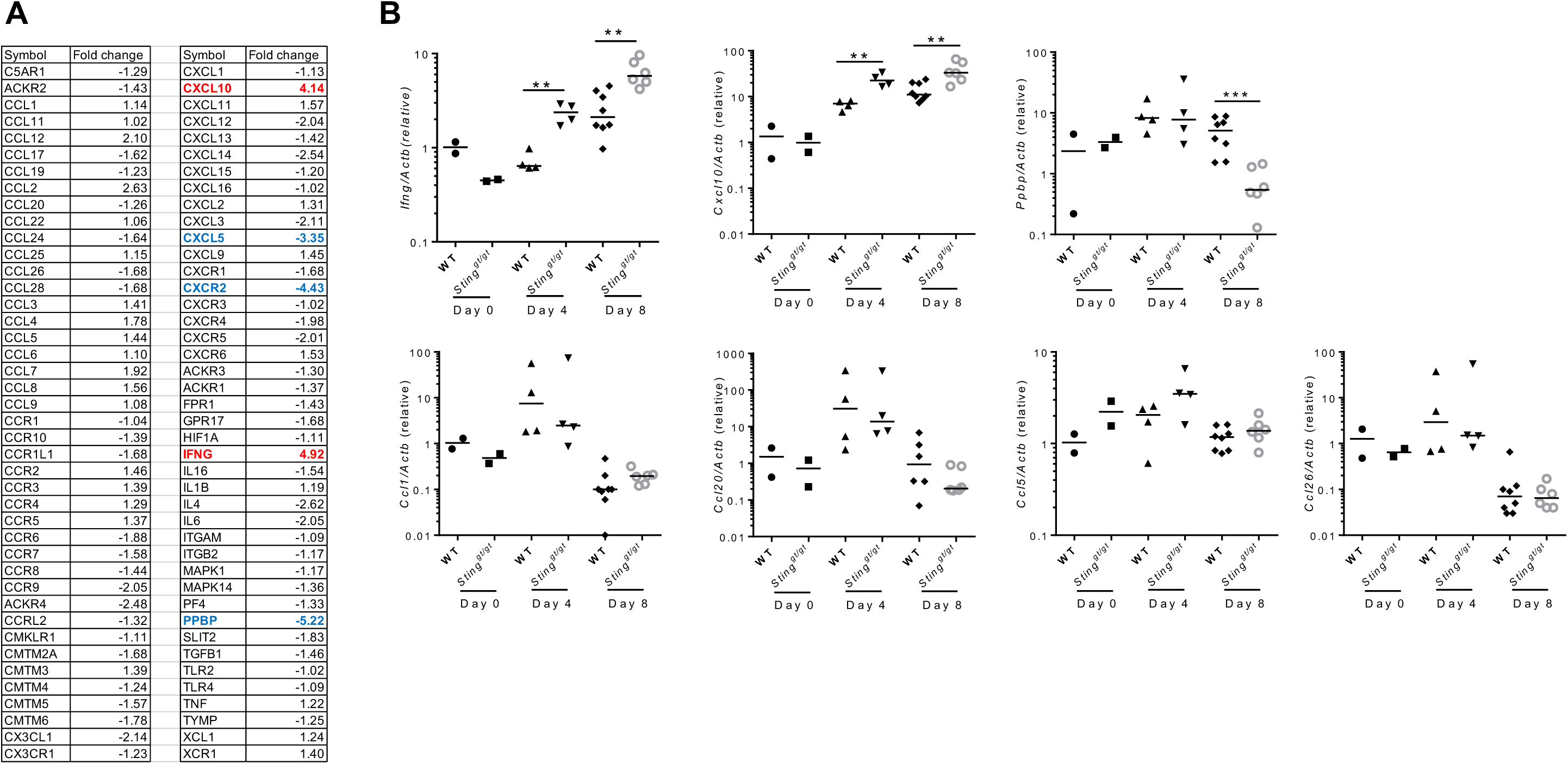
STING deficiency selectively elevates expression of CXCL10 in ankle joints. Age- and sex-matched, C57BL/6 (WT) and STING deficient (*Sting*^gt/gt^) mice were infected with CHIKV. **A**) Multiple genes are differentially expressed in the joints of *Sting*^gt/gt^ (N=6) vs. WT (N=8) mice at day 8 p.i. by PCR array. **B**) qPCR analysis of gene expression at various time points after infection. Each dot=one mouse. The horizontal line in each column=the median. *, P<0.05; **, P<0.01; ***, P<0.001 (non-parametric Mann-Whitney t test).

### CXCL10 signaling contributes to alphavirus pathogenesis

To investigate the physiological role of CXCL10 in alphavirus pathogenesis, we inoculated CHIKV directly into the footpads of both wild-type (WT) and Cxcl10 knockout (*Cxcl10*^-/-^) mice. The results show that the viremia of *Cxcl10*^-/-^ mice were comparable to those in WT mice at days 2 and 4 p.i. (**Fig.3 A**), suggesting that CXCL10 is dispensable for controlling systemic dissemination of CHIKV. The viral loads in the infected feet of WT and *Cxcl10*^-/-^ mice were the same at day 2 p.i., increased modestly in WT at day 4 p.i., while dropped significantly and rapidly in *Cxcl10*^-/-^ mice from days 4 through 7 (**Fig.3 B**), suggesting that CXCL10 signaling promotes viral persistence. Intriguingly, histopathological analyses by H&E staining confirmed a moderate decrease in immune cell numbers in the muscles and joints of *Cxcl10*^-/-^ compared to WT mice (**Fig.3 C**). Consistently, the mRNA expression of *Ifnb1* and inflammatory cytokines (*Il6, Tnfa*) was reduced in *Cxcl10*^*-/-*^ joints at day 7 p.i. compared to WT (**Fig.3 D**). We next asked if this phenomenon is applicable to other arthritogenic viruses. To this end, we chose O’nyong-nyong (ONNV), which together with CHIKV are members of the Semliki Forest antigenic complex of the *Alphavirus* genus. Consistent with the results from CHIKV studies, ONNV viremia were not influenced by *Cxcl10* deficiency (**Fig.4 A**); while the viral loads in the infected feet were lower in *Cxcl10*^-/-^ than those in WT mice at day 6 p.i. (**Fig.4 B**). Histopathological analyses (arbitrary score) by hematoxylin and eosin staining confirmed a moderate decrease in immune cell infiltration into the muscles and joints of *Cxcl10*^-/-^ compared to WT mice (**Fig.4 C**). These data suggest that CXCL10 signaling promotes alphaviral persistence and immune cell infiltration into mouse feet.

**Fig. 3.**
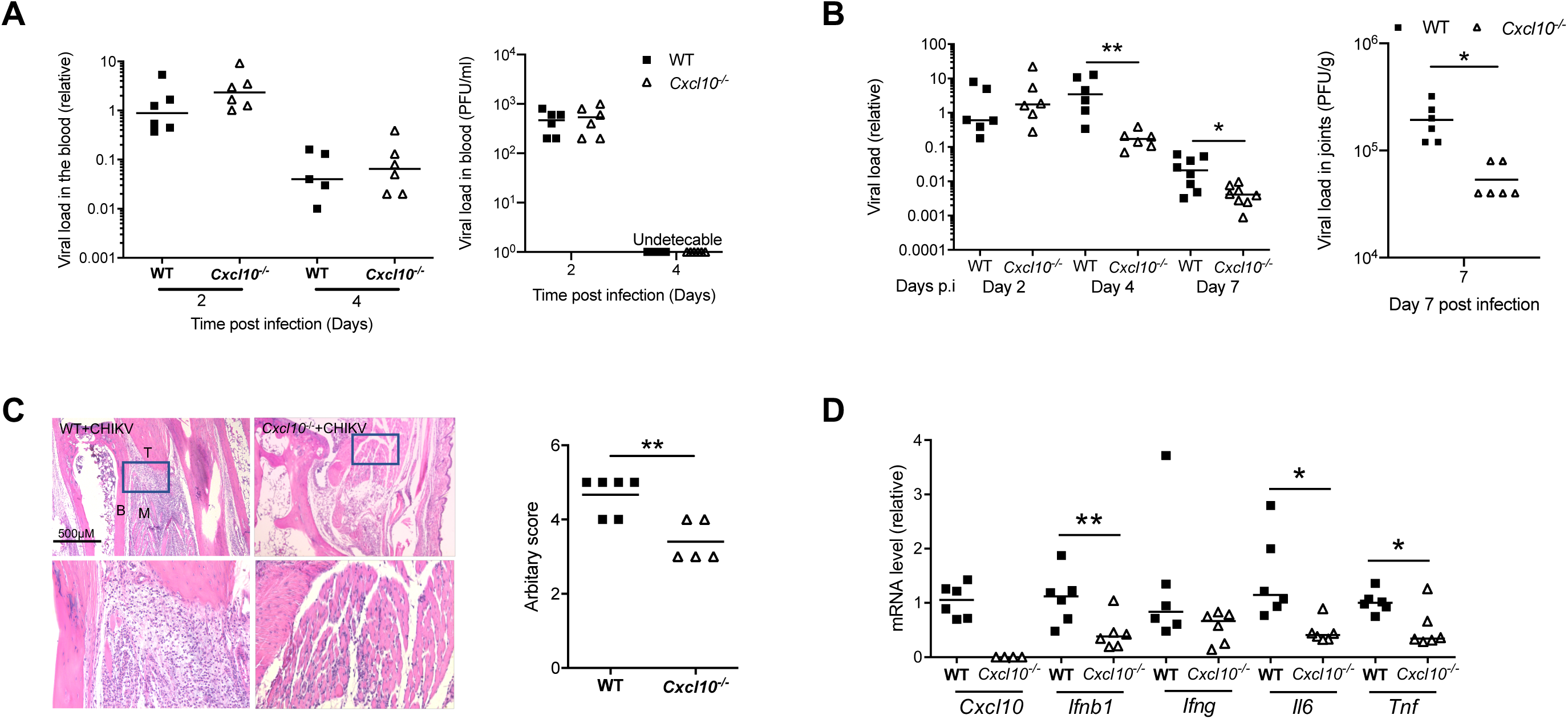
CXCL10 signaling facilitates CHIKV pathogenesis in mouse feet. Age- and sex-matched, C57BL/6 (WT) and Cxcl10 deficient (*Cxcl10*^*-/-*^) mice were infected with CHIKV. **A**) Quantification of viremia by qPCR in whole blood cells (left panel) and plaque assay (PFU/ml serum) (right panel) of wild type (WT) and *Cxcl10*^-/-^ mice. **B**) Quantification of viral loads by qPCR (left panel) and plaque forming assay (PFU/g tissue, day 7) at various days after infection. **C**) Representative H&E micrographs and arbitrary scores of ankle joint inflammation and damage using a scale 1 to 5, with 5 representing the worst disease at 7 days after CHIKV infection. N=5-6/genotype. Magnifications 40X. B: bone, T: tendon, M: muscle. Boxed areas indicate the regions with infiltration and tissue damage. Magnification: 200x. **D**) qPCR quantification of cytokines in ankle joints at day 8 p.i. Each dot=one mouse. The horizontal line in each column=the median. *, P<0.05; **, P<0.01 [(non-parametric Mann-Whitney t test for **B**) and two-tailed Student’s t-test for **D**)].

**Fig. 4.**
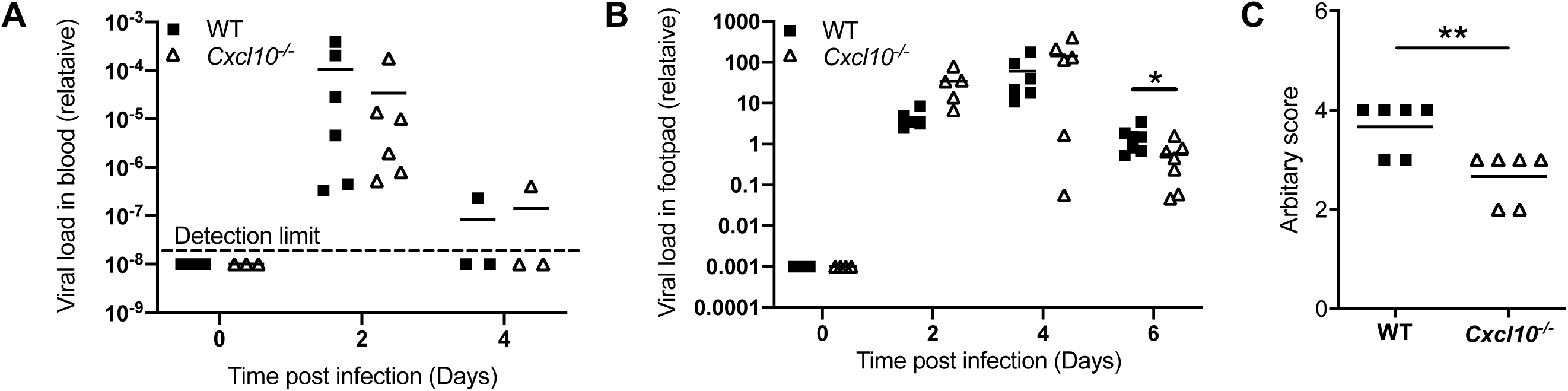
CXCL10 signaling facilitates ONNV pathogenesis in mouse feet. Age- and sex-matched, C57BL/6 (WT) and Cxcl10 deficient (*Cxcl10*^*-/-*^) mice were infected with ONNV. **A**) qPCR quantification of ONNV loads in **A**) whole blood cells and **B**) the ankle joints at various days after infection. **C**) Arbitrary scores of ankle joint inflammation and damage using a scale 1 to 5, with 5 representing the worst disease at 6 days after ONNV infection. N=6/genotype. Each dot=one mouse. The horizontal line in each column=the median. *, P<0.05; **, P<0.01 [(non-parametric Mann-Whitney t test for **B**) and two-tailed Student’s t-test for **C**)].

### CXCL10 signaling promotes macrophage recruitment to infected feet

Since CXCL10 is a chemoattractant for monocytes/macrophages, we analyzed the immune infiltrates in the infected feet by fluorescence activated cell sorting (FACS) to identify and quantitate individual cell populations. In WT mice, total CD45^+^ cells increased modestly at day 2 p.i., decreased a bit at day 4 p.i. and went up again at day 6 p.i. There was a modest decrease in CD45^+^ cells in *Cxcl10*^-/-^ at day 6 p.i. compared to WT (P=0.06) (**Fig.5 A**). Macrophages were recruited rapidly as early as day 2 p.i. and were the largest immune population at all the censored time points. Intriguingly, these cells were significantly fewer in *Cxcl10*^-/-^ than WT mice at day 6 p.i. (**Fig.5 B**). Neutrophils constituted the second largest immune population and infiltrated into the infected feet similarly between WT and *Cxcl10*^-/-^ mice in terms of quantities and kinetics (**Fig.5 C**). Compared to those in the uninfected mice (day 0), the numbers of conventional dendritic cells (cDC) were only significantly increased by day 6 p.i. and higher in *Cxcl10*^-/-^ than WT mice (**Fig.5 D**). Plasmacytoid dendritic cells (pDC) were also recruited to the infected feet as early as day 2 p.i. and their numbers were the same in both genotypes (**Fig.5 E**).

**Fig. 5.**
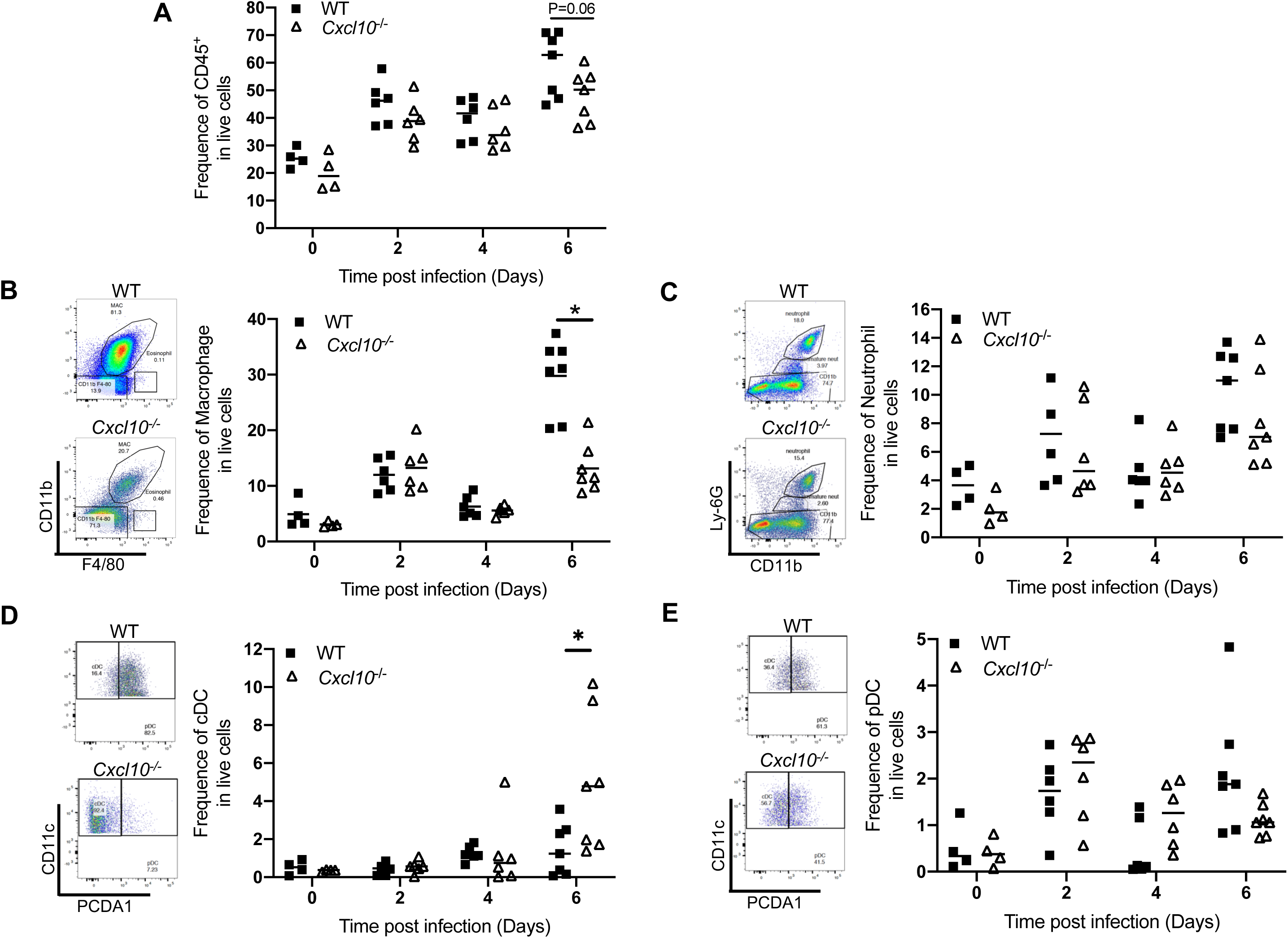
CXCL10 signaling facilitates macrophage infiltration into mouse feet. Age- and sex-matched, C57BL/6 (WT) and Cxcl10 deficient (*Cxcl10*^*-/-*^) mice were infected with ONNV. Different immune cells were quantitated by FACS. The frequencies of **A**) total CD45^+^ immune cells, **B**) CD11b^+^ F4/80^+^ macrophages, **C**) CD11b^+^ Ly-6G^+^ neutrophils, **D**) CD11c^+^ MHCII^+^ conventional dendritic cells (cDC), and **E**) CD11c^+^ PCDA1^+^ plasmacytoid dendritic cells (pDC). Each dot=one mouse. The horizontal line in each column=the median. *, P<0.05 (non-parametric Mann-Whitney t test).

### CXCL10 signaling promotes alphavirus persistence in infiltrating neutrophils and macrophages

The abovementioned data show that macrophages and neutrophils are the primary infiltrating cells in the infected mouse feet, and interestingly the former has been demonstrated to be likely a source of CHIKV persistence in nonhuman primates ^51^. We then examined ONNV RNA loads in each cell population after FACS. We were able to extract RNA from paraformaldehyde-fixed cells using a specialized RNA kit and quantitated viral RNA by qRT-PCR. Among all immune cells, neutrophils contained the highest viral load, which was dramatically reduced in *Cxcl10*^-/-^ mice (P=0.03) compared to WT mice (**Fig.6**). Macrophages had the second highest viral load, which was also decreased in *Cxcl10*^-/-^ (P=0.05). The viral loads in cDC and pDC, though at a much lower level than neutrophil/macrophages, trended lower in *Cxcl10*^-/-^ (**Fig.6**). These data suggest that during the acute phase of infection neutrophils and macrophages are likely an important source of alphaviral replication in infected tissues, and CXCL10 signaling promotes alphavirus persistence in infiltrating neutrophils and macrophages in the foot.

**Fig. 6.**
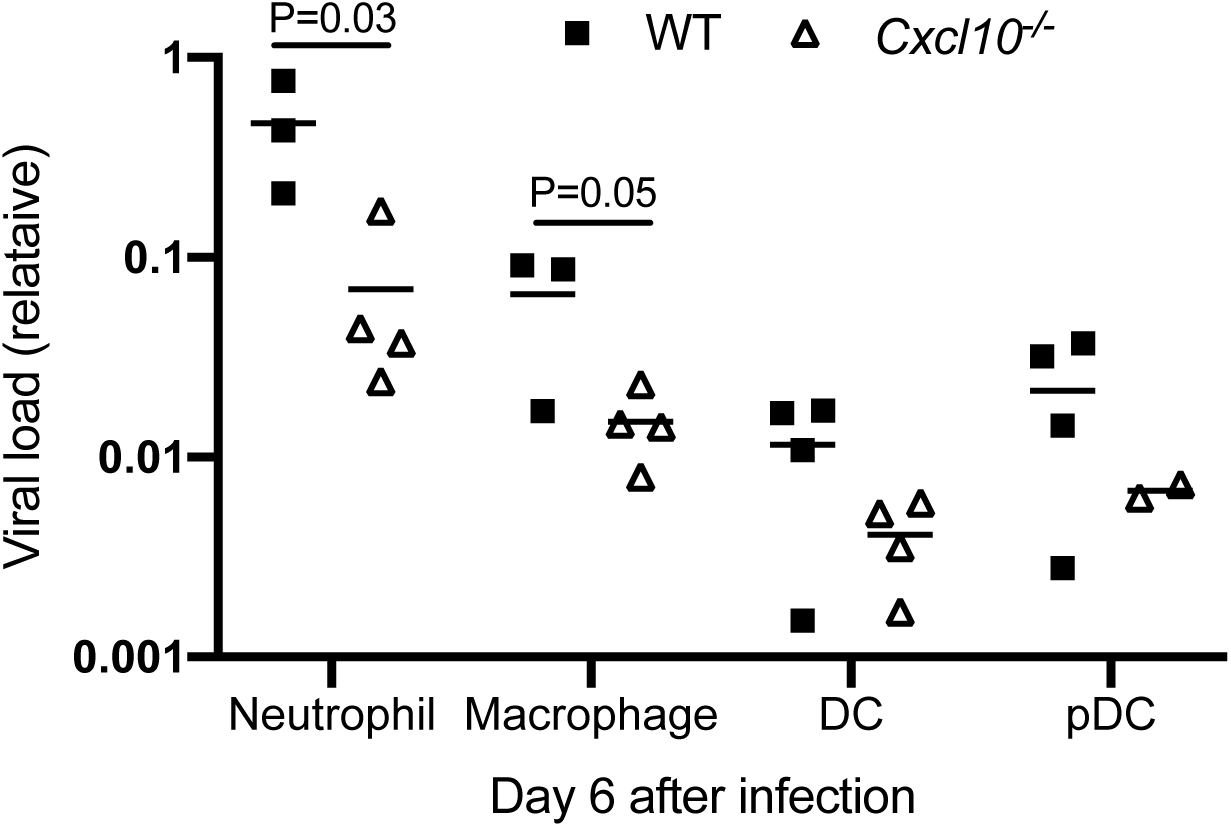
Viral RNA loads are reduced in *Cxcl10*^-/-^ macrophages and neutrophils. Age- and sex-matched, C57BL/6 (WT) and Cxcl10 deficient (*Cxcl10*^*-/-*^) mice were infected with ONNV. Immune cells were sorted by FACS. ONNV RNA in the sorted immune cells was quantitated by RT-PCR. Each dot=one mouse. The horizontal line in each column=the median. P values were calculated with multiple t-tests.

## Discussion

Many studies including ours have firmly established that STING signaling plays a critical anti-RNA virus role ^33,41-43^, likely by multiple mechanism including, but not limited to, induction of type I IFNs ^33 44 45^, a specific set of chemokines via STAT6 ^43^, and translation inhibition of viral gene expression ^46^. In agreement with these published studies, this study shows that STING signaling is also critical for controlling alphavirus infection in mice. Intriguingly, exacerbated pathology progressed in Sting-deficient mouse feet even when viral loads were repressed to similarly low levels between WT and Sting-deficient mice at days 6 and 8 p.i. (**Fig.1 D-H**). These results are in agreement with previous observations showing that CHIKV arthritis severity is not always positively correlated with viral loads ^28^, but rather is a primary consequence of dysregulated inflammatory responses. Thus, our data suggest that STING signaling may not only limit viral replication at the early stage, but also keep aberrant inflammatory responses in check to avoid immunopathology at the later stage. Indeed, STING signaling was recently shown to induce expression of negative regulators of innate immune signaling, such as suppressor of cytokine signaling (SOCS), to control Toll-like receptor (TLR)-mediated hyper inflammatory immune responses in a lupus mouse model ^47^. Plasmacytoid DCs (pDCs) that express a very high level of TLR7 (a viral RNA sensor) was the most rapidly and highly expanded cell type in mouse feet on day 7 after CHIKV/ONNV infection compared to non-infected (**Fig.5 E**) ^52^.

CXCL10 is a chemoattractant for monocytes/macrophages, T cells, NKs, and DCs, and can also promote T cell adhesion to endothelial cells, antitumor activity, and inhibition of bone marrow colony formation and angiogenesis ^53,54,55^. Intriguingly, a high level of serum CXCL10 is associated with severe arthritic disease in humans ^14^. In line with exacerbated joint inflammation, CXCL10 expression was selectively up-regulated in Sting-deficient mice compared to WT (**Fig.2**), suggesting a role for CXCL10 in immunopathology following arthritogenic alphavirus infection. CXCL10 is secreted by several cell types including monocytes, endothelial cells, and fibroblasts. Its expression is increased in many kinds of chronic inflammatory arthritis, especially in rheumatoid arthritis (RA), and is highly induced in CHIKV-infected joints and sustained even after peak viral replication (**Fig.2**). It is thus plausible that during CHIKV infection CXCL10 plays a role in leukocyte homing to inflamed tissues and in perpetuation of inflammation, and therefore, tissue damage. Indeed, joint inflammation was alleviated in *Cxcl10*^-/-^ mice (**Fig.3, 4**), and this was accompanied by a significant reduction in macrophages, which constituted the largest immune cell population in joints following infection (**Fig.5 B**) ^30,51,56^. These activated macrophages could be a main cellular reservoir for CHIKV persistence during the late stages of infection ^30,51^ and contribute to sustained inflammation. In addition to recruiting immune cells, CXCL10 signaling could directly stimulate viral replication, for instance, human immunodeficiency virus 1 (HIV-1) replication in macrophages and lymphocytes ^57^. In this study, we unexpectedly observed a reduction in CHIKV/ONNV in *Cxcl10*^-/-^ compared to WT mouse feet at the late stages (days 4/6 and thereafter respectively) (**Fig.3 B, Fig.4 B**), suggesting a pro-viral role for CXCL10 signaling. However, the absence of CXCL10 did not impact viremia (**Fig.3 A**), suggesting that CXCL10 signaling is dispensable for systemic viral dissemination. Thus, a reduction of viral loads in *Cxcl10*^-/-^ mouse feet could be due to fewer macrophages that are supportive of viral replication in *Cxcl10*^-/-^ than WT feet ^30,51^. Intriguingly, the viral RNA loads in macrophages and neutrophils of *Cxcl10*^-/-^ mouse feet were ∼3-6-fold lower than those of WT mice (**Fig.6**). Macrophages and neutrophils were the predominant immune cell types in mouse feet following alphaviral infection (**Fig.5**). As such, it is plausible that CXCL10 signaling could directly promote alphaviral replication in macrophages and/or other immune cells.

CXCL10 is also a chemoattractant for CD4^+^ T cells, which, though constituting only a small fraction of immune infiltrates during the second peak of foot swelling, are believed to underlie CHIKV-induced inflammation in mice ^16,28,58^. However, CD4^+^ T cells were recruited to footpads by day 6 following ONNV infection and the numbers of CD4^+^ T cells in *Cxcl10*^-/-^ feet were no different than those in WT feet (data not shown).

In summary, our results demonstrate that STING signaling suppresses alphaviral replication and pathogenesis of alphavirus-induced arthritis; while CXCL10 signaling does the opposite. Future work is required to elucidate how CXCL10 signaling facilitates alphaviral replication and test if blockade of CXCL10 signaling mitigates alphaviral arthritis.

## Materials and Methods

### Mice

All the mice used in this study were purchased and bred in our state-of-art animal facility. Wild-type C57BL/6J (JAX Stock #: 000664), *Cxcl10*^-/-^ (JAX Stock #: 000687 on C57BL/6 background) and Sting mutant (*Sting*^gt/gt^ on C57BL/6 background, JAX Stock #: 017537) mice were obtained from the Jackson Laboratory. For each experiment, both sex (both gender)- and age (range 6-12 weeks)-matched WT/mutant mice were used. Mouse experiments were approved and performed according to the guidelines of the Institutional Animal Care and Use Committee at the University of Connecticut and Yale University.

### Cells and Viruses

Vero cells (monkey kidney epithelial cells, Cat. # CCL-81) were purchased from ATCC (Manassas, VA, USA). The cells were grown at 37°C and 5% CO_2_ in complete DMEM medium: Dulbecco’s modified Eagle medium (DMEM) (Corning) supplemented with 10% fetal bovine serum (FBS) (Gibco) and 1% penicillin-streptomycin (P/S; Corning). The CHIKV French La Reunion strain LR2006-OPY1 was a kind gift of The Connecticut Agricultural Experiment Station at New Haven, CT, USA. The ONNV non-recombinant strain was provided by the World Reference Center for Emerging Viruses and Arboviruses (WRCEVA) at University of Texas Medical Branch. Both viruses were propagated in Vero cells.

### Plaque forming assay

Quantification of infectious viral particles in cell culture supernatants/mouse tissue homogenates/mouse sera was performed on a Vero cell monolayer in a 6-well plate following an established protocol ^59^. A serial of 10-fold dilutions of viral samples were prepared in DMEM without fetal bovine serum. In a 6-well plate, 500µL of diluted samples were added to Vero monolayer. The plate was incubated at 37°C and 5% CO_2_ for 2 hrs. The inoculum was then removed and replaced with 2 mL of complete DMEM medium with 1% SeaPlaque agarose (Lonza, Cat# 50100). The plate was incubated at 37°C and 5% CO_2_ for 3 days, and plaques were visualized by a Neutral Red exclusion assay. Viable cells took up neutral red; while dead cells excluded it and thus formed a circular white spot.

### Mouse infection and disease monitoring

Age- and sex-matched mice were inoculated subcutaneously in the hind footpad with 3×10^5^ plaque forming units (PFUs) of CHIKV/ONNV. Mice were monitored for clinical signs of disease afterwards. Footpad swelling was measured using a precision digital caliper.

### Histology studies

Mice were sacrificed and feet were removed and fixed with 4% paraformaldehyde. Tissues were embedded in paraffin and were processed to obtain 5μm sections. Tissues were stained with hematoxylin and eosin. Arthritic disease was arbitrarily scored 1-5, with 5 representing the worst, based on exudation of fibrin and inflammatory cells into the joints, alteration in the thickness of tendons or ligament sheaths, and hypertrophy and hyperplasia of the synovium ^53^. Slides were imaged using an Accu-Scope EXI-310 model inverted microscope with Infinity Capture software.

### Flow cytometry and florescence activated cell sorting

Mice were euthanized, footpads and ankles were harvested at 0, 2, 4 and 6 days post infection (dpi). The footpads were skinned and put into 4 ml of digestion medium with 20 mg/ml collagenase IV (Sigma-Aldrich), 5 U/ml dispase (Stemcell) and 50 mg/ml DNase I mix (Qiagen) in complete RPMI1640 medium. The tissues were harvested and incubated in digestion medium on a shaker at 37°C for 4 hrs. The mixture was transferred to a 40μm cell strainer sitting on a collection tube. 5 ml of complete RPMI medium was added to the cell strainer. Using a circular motion, the digested tissues were ground into the medium against the cell strainer to release maximum number of cells. Cells were then centrifuged at 500xg for 5 min. The supernatant was discarded and red blood cells were lysed using 0.2% sodium chloride. Cells were washed once in complete RPMI medium, re-suspended in 10ml of complete RPMI medium in a 15ml-tube. 10ml of 35% v/v Percoll/RPMI medium was carefully added to the cell suspension. The tube was spun for 20 min at 1200xg. The pellet was re-suspended and washed with complete RPMI medium once.

The isolated cells were then stained for 30 min at 4°C with the following antibodies (Biolegend): APC-Fire 750-anti CD11b (Cat. # 101261), Alexa Flour 700-anti Ly-6G (Cat. # 127621), Brilliant Violet 421-anti CD11c (Cat. # 117343), PerCP-Cy5.5-anti MHC II (Cat. # 107625), PE-anti Tetherin (PCDA1) (Cat. # 12703), Brilliant Violet 510-anti F4/80 (Cat. # 123135), APC-anti CD68 (Cat. # 137007), PE-Dazzle 594-anti CD3 epsilon (Cat. # 100347), Brilliant Violet 711-anti CD4 (Cat. # 100557), Brilliant Violet 570-anti CD8a (Cat. # 100739), FITC-anti CD25 (Cat. # 102005), Zombie UV (Cat. # 423107), PE-Cy7-anti CD45 (Cat. # 103113), TruStain FcX-anti CD16/32 (Cat. # 101319). After staining and washing, the cells were fixed with 4%PFA analyzed by FACS.

Flow cytometry was later performed on a Becton-Dickinson FACS ARIA II, CyAn advanced digital processor (ADP) and analyzed using FlowJo software. Neutrophils were classified as CD11b^+^ Ly6G^+^. Macrophage were classified as CD11b^+^ F4/80^+^. DC cells were classified as CD11c^+^ MHC II^+^. pDC cells were classified as CD11c^+^ PCDA1^+^.

### Real-time quantitative RT-PCR

RNA was isolated from blood samples and footpad tissues using a RNAasy mini-prep kit (Invitrogen). For paraformaldehyde-fixed and sorted cells, RNA was isolated using RNeasy FFPE Kit (Qiagen). Isolated RNA was resuspended in RNAse/DNAse free H_2_O (Invitrogen) and stored at 4°C overnight or −80°C. RT was performed on a Bio-Rad CFX machine using the RNA RT Kit (Takara) with a 10μl total reaction volume per well containing 3μl of RNA samples.

Quantitative PCR (qPCR) was performed with gene specific primers and SYBR Green. The primers for CHIKV were forward primer (5’-GCGAATTCGGCGCAGCACCAAGGACAACTTCA-3’) and reverse primer (5’-AATGCGGCCGCCTAGCAGCATATTAGGCTAAGCAGG-3’). The primers for ONNV were forward primer (5’-GCAGGGAGGCCAGGACAGT-3)’ and reverse primer (5’-GCCCCTTTTTCYTTGAGCCAGTA-3’). The housekeeping gene control used were beta actin, *Actb*. The following PCR cycling program was used: 10 min at 95°, and 40 cycles of 15 sec at 95° and 1 min at 60°C. The results were calculated using the -ΔΔCt method.

### Statistical analysis

All data were analyzed with GraphPad Prism software. For viral RNA analysis, immune cell analysis, cytokines and chemokines analysis and footpad swelling, data were analyzed by the nonparametric Mann-Whitney test, two-tailed Student’s *t* test or multiple t-tests depending on the data distribution and the number of comparison groups. P values of less than 0.05 were considered statistically significant.

## Acknowledgements

We are grateful to The Connecticut Agricultural Experiment Station for providing Chikungunya virus. This work was supported by a National Institutes of Health grant R01AI132526 to P.W.

## Author contributions

T.L. and T.G designed and performed the majority of the experimental procedures and data analyses. A.H. and D.Y. contributed to some of the experiments and/or provided technical support. A.V. and E.F. helped with data interpretations and writing. P.W. conceived and oversaw the study. T.L. and P.W. wrote the paper and all the authors reviewed and/or modified the manuscript.

